# Targeted volumetric single-molecule localization microscopy of defined presynaptic structures in brain sections

**DOI:** 10.1101/568279

**Authors:** Martin Pauli, Mila M. Paul, Sven Proppert, Marzieh Sharifi, Felix Repp, Philip Kollmannsberger, Markus Sauer, Manfred Heckmann, Anna-Leena Sirén

## Abstract

Revealing the molecular organization of anatomically precisely defined brain regions is necessary for the refined understanding of synaptic plasticity. Although, three-dimensional (3D) single-molecule localization microscopy can provide the required molecular resolution, single-molecule imaging more than a few micrometers deep into tissue remains challenging. To quantify presynaptic active zones (AZ) of entire, large, conditional detonator hippocampal mossy fiber (MF) boutons with diameters as large as 10 µm, we developed a method for aberration-free volumetric *direct* stochastic optical reconstruction microscopy (*d*STORM). An optimized protocol for fast repeated axial scanning and efficient sequential labeling of the AZ scaffold Bassoon and membrane bound GFP with Alexa Fluor 647 enables 3D-*d*STORM imaging of 25 µm thick mouse brain sections and assignment of AZs to specific neuronal substructures. Quantitative data analysis revealed large differences in Bassoon cluster size and density for distinct hippocampal regions with largest clusters in MF boutons.

---

Synapses have been imaged in defined functional states with electron microscopy (EM)^1-4^ Serial sectioning^5^ and serial block-face^6^ scanning EM enabled 3D reconstructions of large volumes with high spatial resolution. On the other hand, proteins can be quantified in synapses using freeze fracture EM^3, 7^. Biochemical analysis and super-resolution light microscopy of synaptosomes and hippocampal cultures have led to a 3D model of an “average” synapse^8^. However, since synapses differ substantially from one another even in defined structures e.g. along the dorsal ventral axis of the hippocampus (a key brain area for learning and memory^9^) quantitative information with nanometer spatial resolution in large tissue blocks is highly desirable.

So far single molecule localization microscopy (SMLM) was used successfully to study cultured hippocampal synapses^10-14^. In addition, 3D-SMLM imaging was achieved in brain sections using oil-immersion objectives to visualize the distribution of synaptic proteins a few micrometers above the coverslip^10, 13^. More recently, self-interference 3D super-resolution microscopy and active point-spread function (PSF) shaping in combination with adaptive optics were introduced to enable 3D localization of emitters in tissue with a thickness of up to 50 µm^15, 16^. The latter approach allowed reconstructing super-resolution volumes with an axial depth of several micrometers. However, so far molecular imaging of well-defined larger regions of interest, e.g. all active zones (AZs) in an entire mossy fiber (MF) bouton was not achieved. Both, photobleaching of fluorophores and inefficient labeling of target proteins due to the restricted penetration of antibodies render quantitative molecular imaging, an intrinsic strength of SMLM^14, 17^, in tissue samples complicated.

Here, we focus on large hippocampal mossy fibers boutons (MFBs)^18^ in brain slices. These so-called conditional detonators are complex, structures with tentacle like filopodial extensions and large diameters^5-7^. A single MFB may contain 18 to 45 separate but very closely spaced active zones (AZs) harboring transmitter release sites^5, 6, 19^ and up to 25,000 synaptic vesicles with 1,400–5,700 vesicles as a readily releasable pool^5, 20^. MFBs show remarkable presynaptic plasticity and high variability in presynaptic patch clamp and capacitance measurements^20, 21^, a low release probability (0.01–0.05)^22^ and a loose microdomain coupling^21^. Neither bouton size nor peak calcium current amplitude predict release^20^, therefore it is likely that the molecular organization of AZs controls the plastic nature of MFBs AZs^18, 21, 23^.

To image anatomically precisely defined regions of MFs we developed a standardized sectioning protocol to obtain 1 µm horizontal sections at 2 anatomical levels of the hippocampus: A ventral level 1 and a dorsal level 2 which were 600 µm apart from each other (**Fig. 1a**). Acute 300 µm-thick slices are routinely used for electrophysiological recordings^3, 20-22^ which gives a distance of 600 µm from the ventral to the dorsal surface of 2 successive slices comparable to our 2 levels. To precisely set imaging windows (30 µm x 30 µm) in the middle of the MF tract we used Thy1EGFP(M) mice with strong cytoplasmic expression of EGFP in MFs^24^. The position of the imaging windows in MFs was further confirmed with anti-Zinc-transporter 3 (ZnT-3) staining (**Fig. 1b**). Thin sections are ideal for single-molecule multicolor imaging with red and green absorbing and emitting fluorophores. We used two-color 2D-*d*STORM to resolve the molecular composition of synaptic contacts with antibodies directed against the presynaptic AZ protein Bassoon^25^, and the postsynaptic density protein Homer 1^26^ (**Fig. 1c-e**). Bassoon contains multiple domains for interaction with other AZ components such as RIM-binding protein (RBP), Piccolo and other proteins^25, 27^. Its orientation within the AZ is unclear but it seems to be attached in membrane proximity via its PxxP-motif interaction with RBP (**Fig.1f**). Our commercial monoclonal mouse antibody against Bassoon^10, 25^ has a publically designated epitope covering several hundred amino acids within a so-called Piccolo-Bassoon homology region^25^. We used epitope mapping to identify a highly specific binding site of 9 amino acids from position 875 to 883 (DTAVSGRGL) in the N-terminal region of the molecule (**Fig. 1f**). As Homer 1 is a small protein and is known to form a mesh of heterodimers within the PSD^26^, the C- and N-terminal portions of the molecule should be localized relatively close to each other. Therefore, a more precise epitope mapping of the commercial polyclonal antibody was not necessary (**Fig. 1f**).

**Fig. 1.**
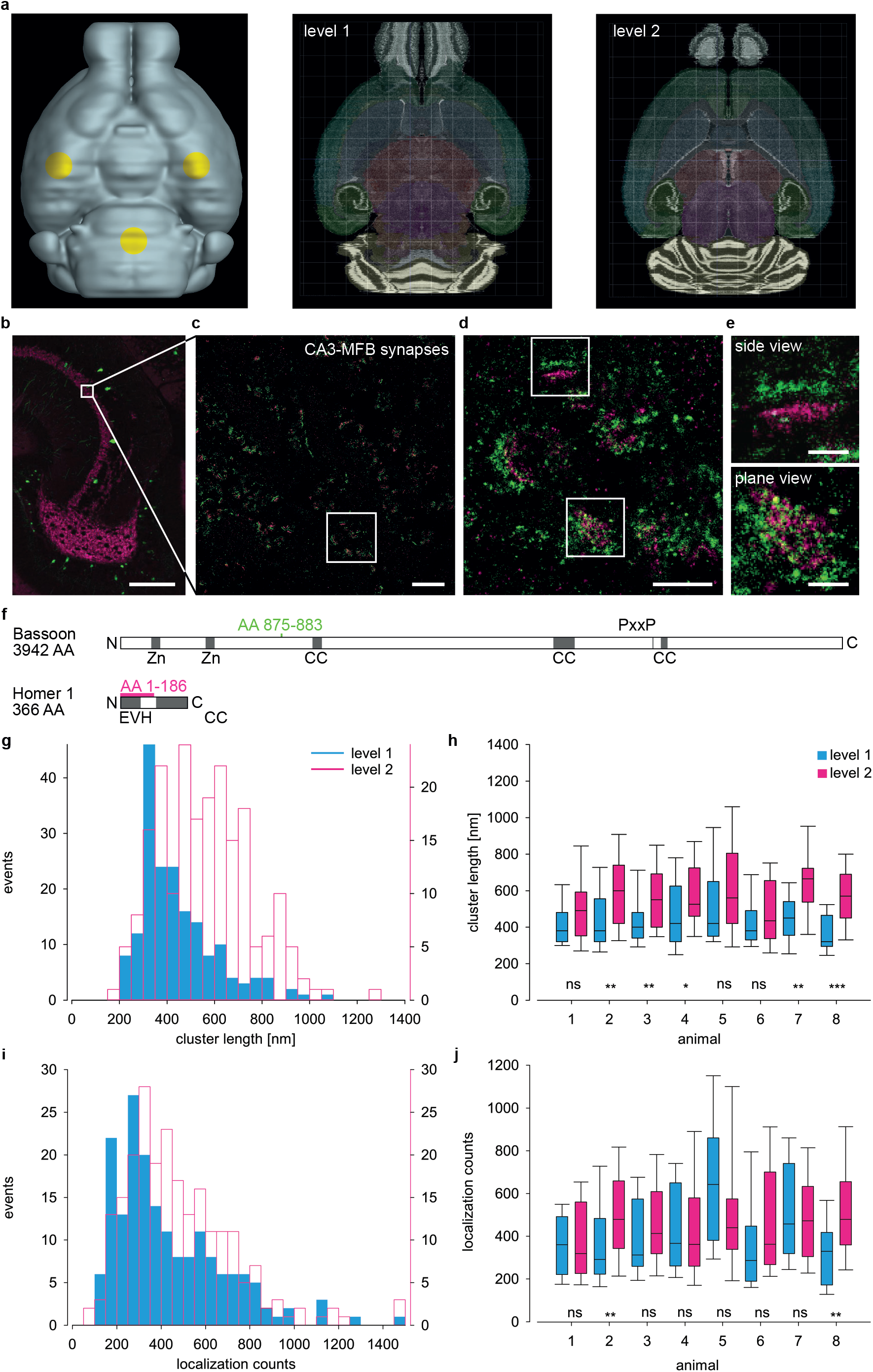
Two-color imaging of hippocampal mossy fibers and Bassoon clusters along the ventro-dorsal axis. (**a**) Ventral view of a virtual mouse brain with landmarks for trimming in yellow (left) and horizontal sections from Allen Mouse Brain Atlas to illustrate the ventral (level 1, middle panel) and the 600 µm more dorsal (level 2) region. (**b**) Confocal image of mossy fibers in a Thy1EGFP(M) mouse with zinc-transporter 3 (magenta) staining. White box marks the imaging window (30 µm x 30 µm) in **c**. (**c**) Two-color *d*STORM image of synapses stained for presynaptic Bassoon (green) and postsynaptic Homer 1 (magenta). White box indicates magnified view in **d**. (**d, e**) Synapses in different orientations: upper box - side view, lower box - plane view further magnified in **e**. (**f**) Protein domain layout of *Mus musculus* Bassoon and Homer1: Zinc-finger (Zn), coiled coil (CC), PxxP and enabled/VASP homology (EVH); epitopes of the anti-Bassoon (green) and anti-Homer 1 (magenta) antibodies. Histograms of length (**g**) and counts (**i**) of Bassoon clusters in side view at level 1 (blue, n = 181 clusters, 9 images, 8 animals) and level 2 (magenta, n = 208 clusters, 7 images, 8 animals). Clusters at level 2 are longer (p < 0.001) and have more counts (p<0.05) than at level 1. Summary box plots (horizontal line median, boxes 25^th^ and 75^th^, whiskers 10^th^ and 90^th^ percentile) in all eight individual animals for length (**h**) and localization counts (**j**) (*p < 0.05; **p < 0.01; ***p < 0.001). (**i**) Scale bars 100 µm in (**b**), 3 µm in (**c**), 1 µm in (**d**) and 200 nm in (**e**).

We identified a large number of contacts in random orientations per image (**Fig. 1c-d**). We focused on those in side views with parallel presynaptic Bassoon (green) and postsynaptic Homer 1 (magenta) (**Fig. 1e**, upper panel). Peak-to-peak distance of Bassoon and Homer 1 clusters was on average 149 ± 12 nm (mean ± SD) showing little variability (**Supplementary Fig. S1a-c**). Clusters in side views were on average 376 ± 136 nm long, had a cross section width of 74 ± 16 nm and 139 ± 80 localization counts per cluster whereas Bassoon clusters in plane view were 460 ± 155 nm long, had a width of 383 ± 98 nm and 297 ± 173 counts per cluster (**Supplementary Fig. S1d-k**). We imaged Bassoon clusters in side views of eight 24-week old male Thy1-EGFP(M) mice (**Fig. 1g-j**). Bassoon cluster lengths and Bassoon localization counts were significantly different at the two levels (**Fig. 1g,i**) and similar differences were obtained when comparing individual animals (**Fig. 1h,j**). Since Bassoon cluster size and neurotransmitter release are correlated^28^ the gradient observed here between level 1 and 2 likely contributes to the high variability observed in recordings from MFBs^20-22^. Furthermore, these differences reinforce the requirement to precisely define regions of interest within mouse brains with submillimeter precision.

To minimize the chance that two clusters are misinterpreted as one in 2D projections, we implemented 3D-*d*STORM. Whereas the xy-view shows a Bassoon cluster that appears to extend to the left forming one large cluster (**Fig. 2a**) 3D images (xz- and yz-view) (**Fig. 2b,c**) clearly show two near-by clusters (**Supplementary Video 1**). A limitation of these images is that many clusters extend beyond the borders of our 1 µm thin brain sections and are thus truncated (**Fig. 2d-h**) as demonstrated in an example of an entire cluster within the slice (**Fig. 2e**) and a truncated one at the upper border of the brain slice (**Fig. 2g**) and in the histograms of the clusters in seven images from one animal (**Fig. 2h-i**). Re-evaluation of the data filtered for truncated clusters reduced cluster volume and variability (median ± 25^th^-75^th^ percentile 0.0078 ± 0.0018-0.0145 µm^3^, n = 1933 in filtered vs. median 0.0102 ± 0.0034-0.0240 µm^3^, n = 3783 in not filtered data) (**Fig. 2h-i**). Thin sections thus may lead to underestimations of cluster size.

**Fig. 2.**
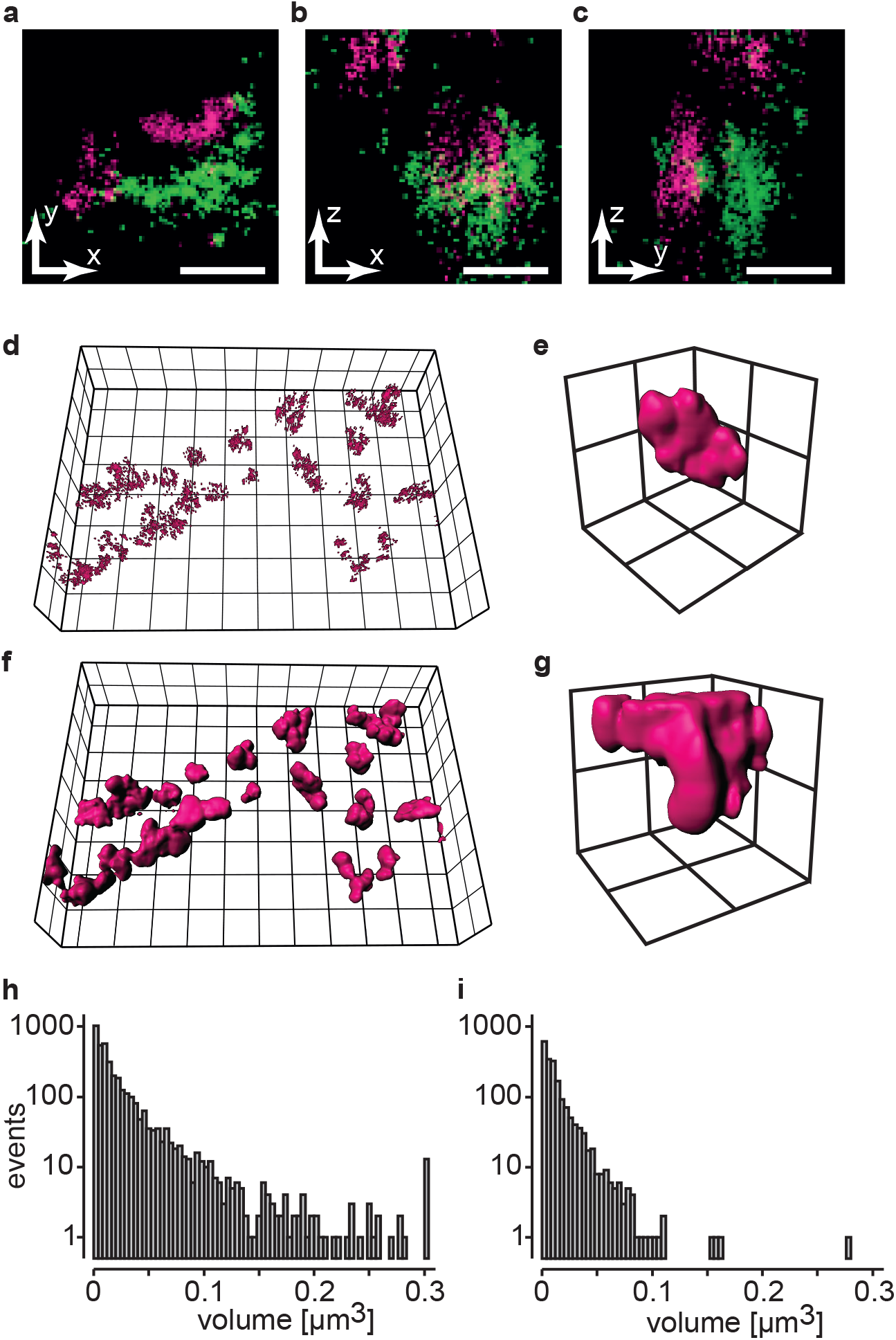
3D imaging uncovers truncation of Bassoon clusters. (**a**-**c**) Representative two-color 3D *d*STORM images of an individual synaptic contact with presynaptic Bassoon (green) and postsynaptic Homer 1 (magenta) in all three orientations (xy-, xz- and yz-view; see **Suppl. Video 1**). (**d**) A localization dataset of Bassoon (magenta) and (**f**) its volume reconstruction. An entire cluster (**e**) within the imaged tissue volume, and a truncated cluster (**g**) extending beyond its upper edge. Semi-logarithmic histograms of cluster volumes in seven images from 1 animal for all clusters (**h,** n = 3783) and for selected non-truncated clusters (**i**, n = 1869). Scale bars in (**a**-**c**) 100 nm, grid size in (**d**) and (**f**) 2 µm and in (**e**) and (**g**) 500 nm.

To reduce truncation, we imaged larger tissue volumes *en bloc* in 25 µm sections using optimized procedures for homogenous labeling (**Methods and Supplementary Fig. S2a**). We imaged three different regions (**Fig. 3a**), MF tract (MFT), perforant path (PP) and Schaffer collaterals (SC) using Thy-1 EGFP fluorescence as a marker for hippocampal architecture in the same tissue slice in less than 3 hours. This enabled an unbiased evaluation of clusters with identical staining conditions for all regions without inter-animal variability. We observed different distributions of clusters in MFT (**Fig. 3b**), PP (**Fig. 3c**) and SC (**Fig. 3d**) with smallest and most sparse clusters in SC. Bassoon localization counts (median ± 25^th^-75^th^ percentile: 22 ± 3-48, n=11232), cluster length (0.291 ± 0.223-0.426 µm) and volume (0.0033 ± 0.0019-0.0074 µm^3^) in MFT were significantly larger (p<0.001) compared to PP (17 ± 12-30; 0.257 ± 0.214-0.336 µm; 0.0026 ± 0.0018-0.0044 µm^3^, n=12596) and SC (16 ± 12-25; 0.248 ± 0.210-0.314 µm; 0.0024 ± 0.0017-0.0038 µm^3^, n=8669) (**Fig. 3e-g**).

**Fig. 3.**
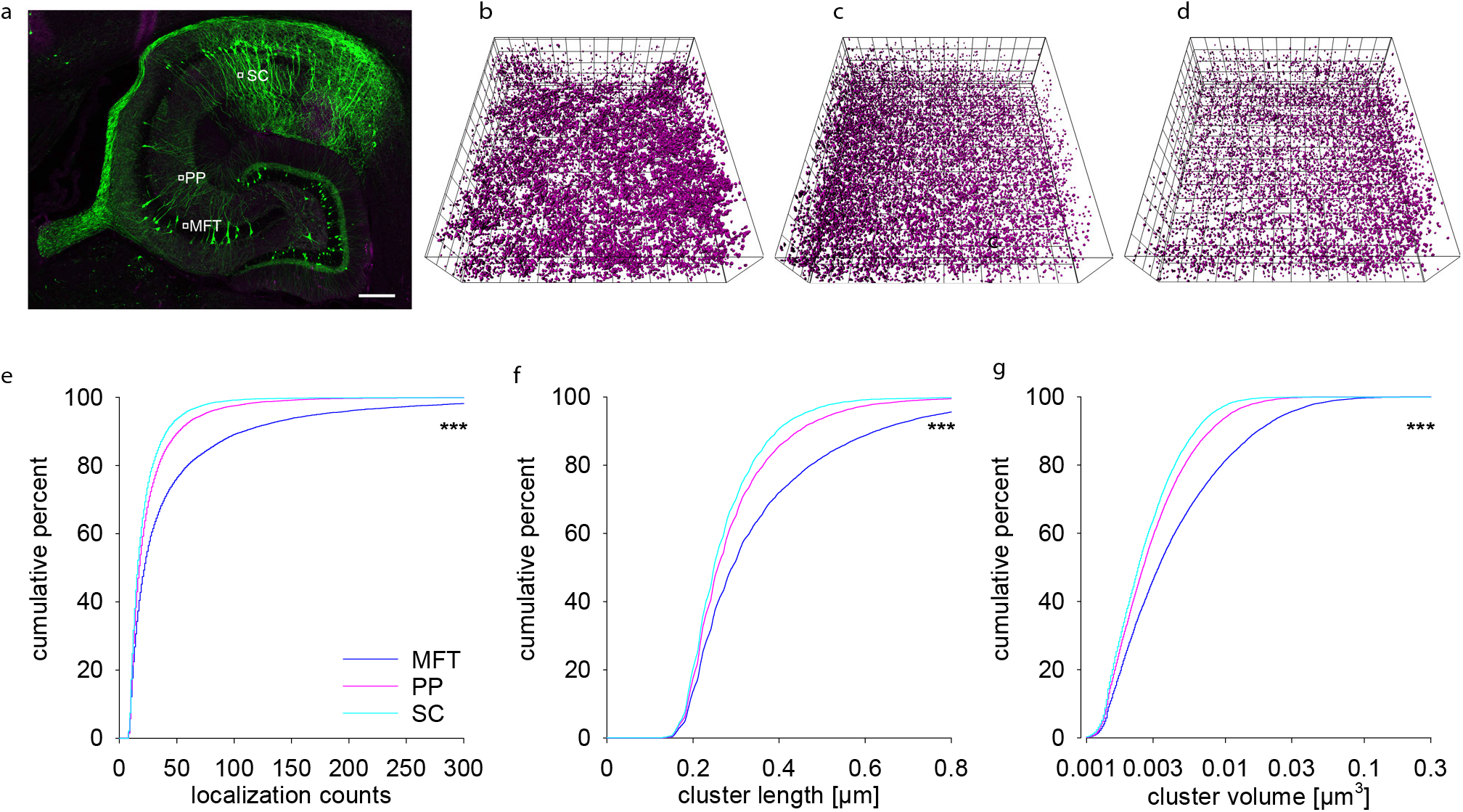
*En bloc* 3D imaging in 25 µm thick tissue slice reveals different Bassoon patterns in three distinct hippocampal circuits. (**a**) Confocal overview of a 25 µm thick section of the hippocampus in a Thy-1 EGFP(M) mouse with imaging windows in the mossy fiber tract (MFT), perforant pathway (PP) and Schaffer collaterals (SC) depicted to scale (25 × 25 µm). 3D plots of Bassoon clusters in MFT (**b**), PP (**c**) and SC (**d**). Cumulative plots of localization counts (**e**), cluster length (**f**) and volumes (**g**) in MFT (dark blue), PP (magenta) and SC (cyan). (**c**-**e**) Scale bar: 200 µm in (**a**) and grid size 2 µm in (**b**-**d**). Asterisks (*** p<0.001) denote statistical significance MFT vs PP/SC and PP vs SC.

For visualization of bouton surfaces we turned to Thy1-mEGFP(Ls1) mice expressing membrane bound EGFP^29^ in which large MFBs were clearly visible as shown in the confocal overview of the hippocampal dentate gyrus (**Fig. 4a**). In the 3D-*d*STORM reconstruction of the anti-GFP fluorescence (**Fig. 4b, Supplementary Video 2**) MF boutons, filopodia and axons are illustrated. We used sequential imaging to map Bassoon clusters to identified large MFBs. Brain slices were mounted on fluorescent bead-coated coverslips which allowed to align sequential images and to correct for drift (**Supplementary Fig. S2**). We first visualized Bassoon with the monoclonal mouse anti-Bassoon and Alexa647-conjugated anti-mouse secondary antibodies. After washing we stained the tissue with Alexa647-conjugated anti-GFP nanobodies (**Supplementary Video 2**). Repeated axial scanning (10 times) in the middle of the 25 µm thick brain slice through the 10 µm thick region of interest in a zig-zag manner showed homogeneous distributions of localization intensity and counts (**Fig. 4c and Supplementary Fig. S2b-c**). Localization counts, cluster length and volume of Bassoon clusters that could be assigned to MFBs (median ± 25^th^-75^th^ percentile: 36 ± 14-96 counts; length 0.558 ± 0.352-0.848 µm, volume 0.0144 ± 0.0052-0.0380 µm^3^, n=181) were larger (p<0.001) than those of Bassoon clusters in the whole imaging window (counts: 22 ± 12-49; length 0.433 ± 0.315-0.629 µm; volume 0.0079 ± 0.0040-0.0193 µm^3^, n=35632) (**Fig. 4d-f**). 21 individually identified boutons showed different volumes and number of clusters per bouton with on average 9 clusters per large MFB (**Supplementary Table 1**). Sequential *d*STORM-imaging thus enabled volumetric protein cluster mapping in identified subcellular structures.

**Fig. 4.**
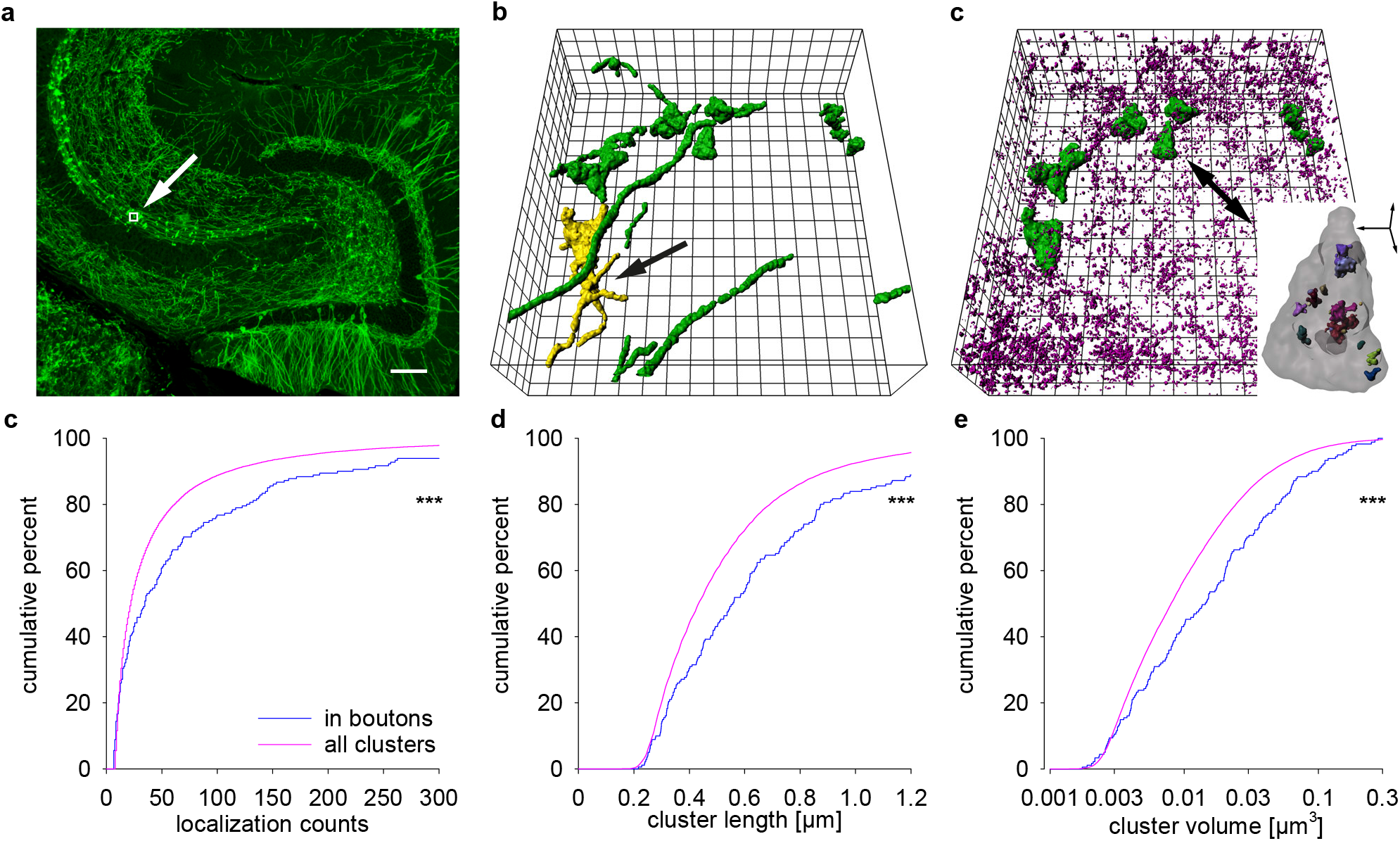
Sequential *en bloc* 3D imaging in 25 µm thick tissue slices of mEGFP(Ls1) mice shows largest Bassoon clusters in identified mossy fiber boutons. (**a**) Confocal overview of mossy fibers in a Thy-1 mEGFP(Ls1) mouse with imaging window (25×25 µm) depicted to scale. (**b**) 3D volume reconstruction of mEGFP in xyz-view, showing axons and mossy fiber boutons (green), a typical bouton with filopodial extensions is marked in yellow (black arrow). (**c**) 3D reconstruction of Bassoon (magenta) and mEGFP (green) in large mossy fiber boutons used for calculations of “clusters in boutons” in (**d**-**f**); inset depicts the marked bouton (black arrow) with Bassoon clusters (colored). Cumulative plots of localization counts (**d**), cluster length (**e**) and volumes (**f**) comparing all clusters per image (magenta) and those in identified boutons (dark blue). Scale bar: 100 µm in (**a**), grid size 2 µm in (**b-c**) and scale bar in inset in (**c**) 1µm. Asterisks (*** p<0.001) denote statistical significance between the groups.

To conclude, we imaged a 30 × 30 µm field of view allowing us to measure typically more than 20 synaptic contacts within a single focal plane by 3D-*d*STORM. Using water immersion, astigmatic imaging with the red fluorophore Alexa Fluor 647 and fast repeated axial 3D scanning at total acquisition times of 3 h resulted in homogeneous localization intensity and counts. Drift stabilization was optimized by automated remote scanning after sample stabilization and correction by imaging beads at the beginning and end of each scan. We took great care to address a number of potential confounding factors such as projection artifacts in 2D-imaging and mechanical or optical sectioning artifacts, which could be excluded as factors accounting for the observed variability. The large size and highly complex geometry of MFBs with on average less than 0.5 µm from one synaptic contact to the next in EM^5, 6, 19^ represents a major imaging challenge further complicated by the highly variable size of individual contacts. The size of the individual clusters in 3D images of Bassoon falls within the range expected for AZs based on previous EM work^5, 7, 19^. Our previous attempts to obtain accurate measurements of Bassoon cluster size for MFBs using confocal, structured illumination microscopy (SIM) or stimulated emission depletion (STED)-microscopy failed due to insufficient z-axis resolution (unpublished data). In particular, the smallest and the largest clusters, which contribute significantly to the overall variability, are challenging. Here, we could characterize the geometry and size of protein clusters in neighboring synaptic circuits within one single anatomically preserved mouse hippocampus section avoiding cutting and tissue processing artifacts. Furthermore, with our method and sequential staining it was possible to image two proteins with Alexa Fluor 647, the most reliable *d*STORM dye for quantitative imaging^17, 30^. Using sequential staining and *en bloc* 3D imaging we were able to characterize the geometry and size of protein clusters and also map clusters to identified MFBs. The clusters in MFBs were significantly larger than the clusters recorded in the whole region of interest and with mice expressing membrane bound EGFP it was possible to distinguish between contacts formed by the MFB itself and its GABAergic filopodial extensions. Further presynaptic structures, e.g. from interneurons likely contributed to the smaller clusters in our images^31^.

The approach described here can be used to map the 3D distribution of any protein in thick tissue slices for which highly specific antibodies are available. While we used mice expressing membrane bound EGFP to identify MFBs the described method can likewise be modified to correlate function and molecular organization of individual synapses in acute slices or even *in vivo* by filling of electrophysiologically characterized synapses with fluorescent dyes for identification and subsequent labeling and volumetric 3D-*d*STORM imaging.

## Materials and methods

### Animals

One 12-week old and eight 24-week old male Thy1-EGFP(M) mice on C57/BL6 background and one 12-week old male C57/BL6 were used for Fig 1-3. Three 12-week old male Thy1-mEGFP(Ls1) mice^29^ with membrane bound mEGFP in hippocampal mossy fiber boutons (MFB) were used for Fig. 4. All animal procedures complied with the guidelines of institutional and regulatory authorities and were conducted in accordance with the EU Directive 2010/63/EU, and the United States Public Health Service’s Policy on Humane Care and Use of Laboratory Animals.

### Cryosectioning

The mice were sacrificed with CO_2_ and perfused transcardially with phosphate buffered saline (PBS) followed with 4% paraformaldehyde (PFA) in 0.2 M PBS. Brains were removed, post-fixed in 4% PFA overnight at 4°C, dehydrated in 30 % sucrose solution and rapidly frozen. 1 µm thin horizontal cryosections were obtained using a cryotome (Leica CM 3050, Leica, Wetzlar, Germany) in a standardized manner. To obtain a uniform cutting angle for horizontal brain sections, brains were mounted with the dorsal cortical surface facing down on aluminum carriers. Using the lower edges of both cortices and the pons as landmarks to align the specimen to the blade, the brains were trimmed until lateral ventricles were opened and the characteristic double C shape of the hippocampus could first be identified at both sides. From this base level 100 µm of the brain were cut in 1 µm thin slices on silan coated coverslips and the following 100 µm were cut in 10 µm thick sections on regular microscope slides. To align the cutting level between different animals on the identical hippocampal structure we used images from Nissl stained horizontal brain atlas (brainmaps.org)^32^ and aligned the sections according to the shape of the dentate gyrus. Cryosections between 900 µm and 1000 µm corresponding to the image 141 of the brain atlas are referred to as level 1 and and sections between 1500 µm and 1600 µm corresponding to image 117 of the atlas as level 2. For *en bloc* 3D *d*STORM imaging, horizontal 25 µm thick cryosections were cut into 12-well tissue culture plates for staining free-floating and mounted after the last washing step on fluorescent beads coated coverslips (see below).

### Silan coated coverslips

Round shaped, 18 mm diameter, high precision coverslips (Marienfeld No 1.5H, Lauda-Königshofen, Germany) were mounted vertically in a custom-made comb-like PTFE-holder, immersed with 3-aminopropyl-triethoxysilane (Sigma 440140, Sigma-Aldrich, Schnelldorf, Germany) 2% v/v in methanol for two minutes, washed shortly with methanol 100% and distilled water and dried. For sequential *d*STORM imaging coverslips were coated with FluoSpheres®, 0.2 µm, orange fluorescent (540/560) (Thermo Fisher Scientific, Darmstadt, Germany, dilution 1:1000). FluoSpheres were diluted in PBT 0.125%. 17 µl were spread on silan-coated coverslip over an area of approximately 1.6 cm^2^, dried, washed with destilled water, and dried again.

### Immunofluorescence

For immunofluorescence staining of 1 µm sections on coverslips, the sections were washed with 0.02 M glycine (Sigma) in PBS for 30 min, blocked with blocking solution consisting of 1% BSA (Sigma) and 5 % NGS (Seralab, West Sussex, UK) in 0.3 % PBT (PBS containing 0.3 % Triton X-100, Sigma) for 90 min and incubated with primary antibodies at 4°C overnight. Samples were washed with blocking solution twice for 5 and twice for 20 minutes, followed by incubation with secondary antibodies for 2 hours at room temperature. After identical washing steps, sections were kept in 1xPBS at 4°C until imaging. The staining procedure for free floating 25 µm sections was identical to that for 1 µm sections, except that the blocking was done overnight, primary antibody incubation for 36 h and secondary antibody incubation for 24 h. After the last washing step, the sections were mounted on beads-covered coverslips, quickly dried, and kept in 1xPBS at 4°C until imaging. Primary antibodies were used in the following concentrations: mouse monoclonal antibody (mAb) anti-α-Bassoon (Enzo Sap7F407, Enzo Life Sciences, Lörrach, Germany, 1:500), rabbit polyclonal antibody anti-α-Homer 1 (Synaptic Systems, Göttingen, Germany, 106 002, 1:500), rabbit polyclonal and mAb anti-Zinc transporter 3 (ZnT3) (Synaptic Systems 197 002 and 197 011, 1:500). Secondary antibodies were used in the following concentrations: Alexa 647 conjugated goat α-mouse Fab, Alexa 532 conjugated goat α-mouse Fab, Alexa 647 conjugated goat α-rabbit Fab, Alexa 532 conjugated goat α-rabbit Fab (Invitrogen, Thermo Fisher, Darmstadt, Germany,1:500 each).

PEPperMAP® Type 1 Epitope Mapping of the mouse monoclonal anti-Bassoon antibody against two undisclosed antigens was performed with 15 amino acid antigen-derived peptides with a peptide-peptide overlap of 14 amino acids by PEPperPRINT GmbH (Heidelberg, Germany). The antigen-derived peptide microarrays were incubated with the antibody samples at concentrations of 1 µg/mL and 10 µg/mL in incubation buffer followed by staining with the secondary antibodies goat anti-mouse IgG (H+L) DyLight680 or sheep anti-rabbit IgG (H+L) DyLight680 and read-out with a LI-COR Odyssey Imaging System. Quantification of spot intensities and peptide annotation were done with PepSlide® Analyzer (PEPperPRINT).

### *d*STORM (*direct* stochastic optical reconstruction microscopy)

Cover slips with stained brain slices were mounted in a custom-made holder and incubated in imaging buffer (100 mM mercaptoethylamine (MEA, Sigma) in PBS, buffered at pH 7.8-7.9) to allow reversible switching of single fluorophores during data acquisition. Images were acquired using an inverted microscope (Olympus IX-71, Olympus, Hamburg, Germany, 60x, NA 1.45, oil immersion) equipped with a nosepiece-stage (IX2-NPS, Olympus). 644 nm (iBEAM-SMART-640-S, Toptica, Gräfelfing, Germany) and 532 nm (Qioptiq Nano 250-532, Qioptiq; Asslar, Germany) lasers were used for excitation of Alexa Fluor 647 and Alexa Fluor 532, respectively. Laser beams were passed through a clean-up filter (Brightline HC 642/10, Semrock, and ZET 532/10, Chroma, AHF Analysetechnik, Tübingen, Germany, respectively) and two dichroic mirrors (Laser-MUX BS 514-543 and HC-quadband BP, Semrock) onto the sample. The emitted fluorescence was filtered with a quadband-filter (HC-quadband 446/523/600/677, Semrock) and divided onto two cameras (iXon Ultra DU-897-U, Andor, Acal BFi, Grübenzell, Germany) using a dichroic mirror (HC-BS 640 imaging, Semrock). In addition, fluorescence was filtered using a longpass- (Edge Basic 635, Semrock) or bandpass-filter (Brightline HC 582/75, Semrock) for red and green channels, respectively. Pixel sizes were 126 nm (red) and 128 nm (green). Single fluorophores were localized and high resolution-images were reconstructed with *rapid*STORM (10 nm /pixel sub-pixel binning)^33^ (www.super-resolution.de).

### 3D-dSTORM

3D-*dS*TORM images were obtained at a setup mainly as described above with a Piezo z-stage (P-736.ZR 2, Physik Instrumente, Karlsruhe, Germany) and a cylindrical lens in the emission path to obtain information about the three dimensional position of the fluorophore^34^. For z-calibration of the microscope, we used multi-fluorescent beads (100 nm TetraSpeck, ThermoFisher) adsorbed on a coverslip and covered with water. The calibration sample was axially moved at constant speed through the focal plane by the piezo and the widths of the PSF in x and y is evaluated with *rapid*STORM 3.3.1^33^ (www.super-resolution.de). The interpolations of the widths against the known z position is performed using cubic B-splines and serve as calibration table^35^. Axial position of localizations in samples was determined using *rapid*STORM as previously described ^35^.

### Sequential en bloc 3D *d*STORM

For scanning sequential 3D-*d*STORM a 647 nm Laser (F-04306-113, MPB Communications Inc., Pointe-Claire, Quebec, Canada) was used. For imaging Alexa647 with long term pH stability at pH 7.8 we used reducing agent 2-mercaptoethylamine-hydrochloride (MEA, 100 mM, Sigma) in a 0.2 M sodium phosphate buffer and a oxygen-scavenging system (10% (wt/vol) glucose, 10 U/ml glucose oxidase and 200 U/ml catalase. Samples on coverslips with fluorescent beads were placed in a custom build imaging chamber and mounted on the microscope with 14 µl of immersion media (Immersol W (2010), Zeiss, Jena, Germany). To minimize drift measurements were started via remote control from outside the laboratory 45 min after the region of interest was defined. Each measurement consisted of several videos with 15,000 frames at 100 Hz each. In order to reconstruct larger volumes we used the piezo for continuous axial scanning over the whole z-range of the ROI during one movie. To counter effects of out of focus photobleaching we performed repeated scanning: typically 10 scans were performed inverting the direction of the movement after each scan in a zig-zag manner. The measurement sequence was controlled by a custom-built micromanager^36^ plugin. For sequential imaging samples were washed with PBS and the second staining procedure was performed as described above. Before and after each measurement fluorescent beads at the surface of the coverslip were imaged for alignment of sequential scans and drift correction. Precise position of beads was determined with *rapid*STORM 3.3.1 and alignment of sequential stacks was performed using elastic transformations obtained with ImageJ^37^ plugin bunwarpJ^38^.

### Data evaluation

Raw localization data obtained from *rapid*STORM 3.3.1 were examined and further processed with FIJI ^37^ (Fig. 1-2) and Imaris (Bitplane, Zürich, Switzerland) (Fig. 3-4).

### 2D imaging

To reflect the complex shape of Bassoon clusters, area measurements were performed using the freehand ROI tool, length and width measurement were performed using the freehand line respectively line tool in FIJI. To obtain an estimate for relative protein content within an AZ we measured localization counts per AZ, i.e. the total number of stochastic fluorophore blinking events per cluster. Using standardized imaging conditions the number of localization counts depends on the number fluorophores present at the target structure, which in turn depends on the number of bound secondary and primary antibody and thereby on the number of epitopes^17^.

### Data processing and analysis for sequential en bloc 3D *d*STORM

We used custom Python 2.7 scripts to prepare the *rapid*STORM localization files for further processing with Imaris (Bitplane). Localization tables from individual scans were concatenated and true z position of each localization was calculated using information about the z-position of the piezo during scanning. Drift correction was applied using rigid transformations obtained from bead recordings before and after each measurement. Finally, images were filtered using density-based clustering (DBSCAN) in sckit-learn^39^ to remove noise. A cut-off of 8 localisations was used for filtered data (see Histogramm in **Supplemental Fig. S3**). After individually processing both channels in sequential scans both channels were combined and loaded into Imaris via the “Super Resolution Localization Data To Image – XTension” and the “Super Resolution Localization Data To Spots – XTension”.

Once loaded, we used the Imaris surface module for volume data extraction of Bassoon clusters and reconstruction of mossy fiber boutons. Boutons were separated manually from supporting axons. Identification of Bassoon clusters belonging to one bouton was done using Imaris XTension distance transformation. All Bassoon clusters with a center of mass inside a reconstructed super-resolved bouton signal were considered to belong to this bouton.

### Statistics

Statistical analyses were performed with Sigma Plot 12 and 14 (Systat Software GmbH, Ekrath, Germany) using the non-parametric Mann-Whitney rank sum test or the non-parametric ANOVA for multiple comparisons. Asterisks indicate the significance level (* p < 0.05, ** p < 0.01, *** p < 0.001). Data are reported as median ± 25^th^ and 75^th^ percentile for non-parametric data unless indicated otherwise and as mean ± SD for parametric data.

## Acknowledgements

The authors thank L. Behringer-Pliess for labelling the Alexa Fluor 647 conjugated nanobodies and T. Trnetschek and B. Gado for further technical assistance. This work has been supported by the German Research Foundation (TRR 166 ReceptorLight, projects B06 and A04), and the Interdisciplinary Clinical Research Center (IZKF) Würzburg (N-229).

## Author contributions

M.P., M.M.P., M.H. and A.-L.S. designed the study, M.P. together with M.M.P., M.SH., S.P. and A.-L-S. performed the experiments and analyzed data together with F.R. and P.K.. A.-L.S., M.H., and M.S. supervised the project and provided funding. M.P. together with A.-L.S. M.H., M.M.P. and M.S. wrote the manuscript. All authors read and approved the manuscript.

