## Supplemental Figures and Table for "Targeted volumetric single-molecule localization microscopy of defined presynaptic structures in brain sections"

### Supplemental figures and table with legends

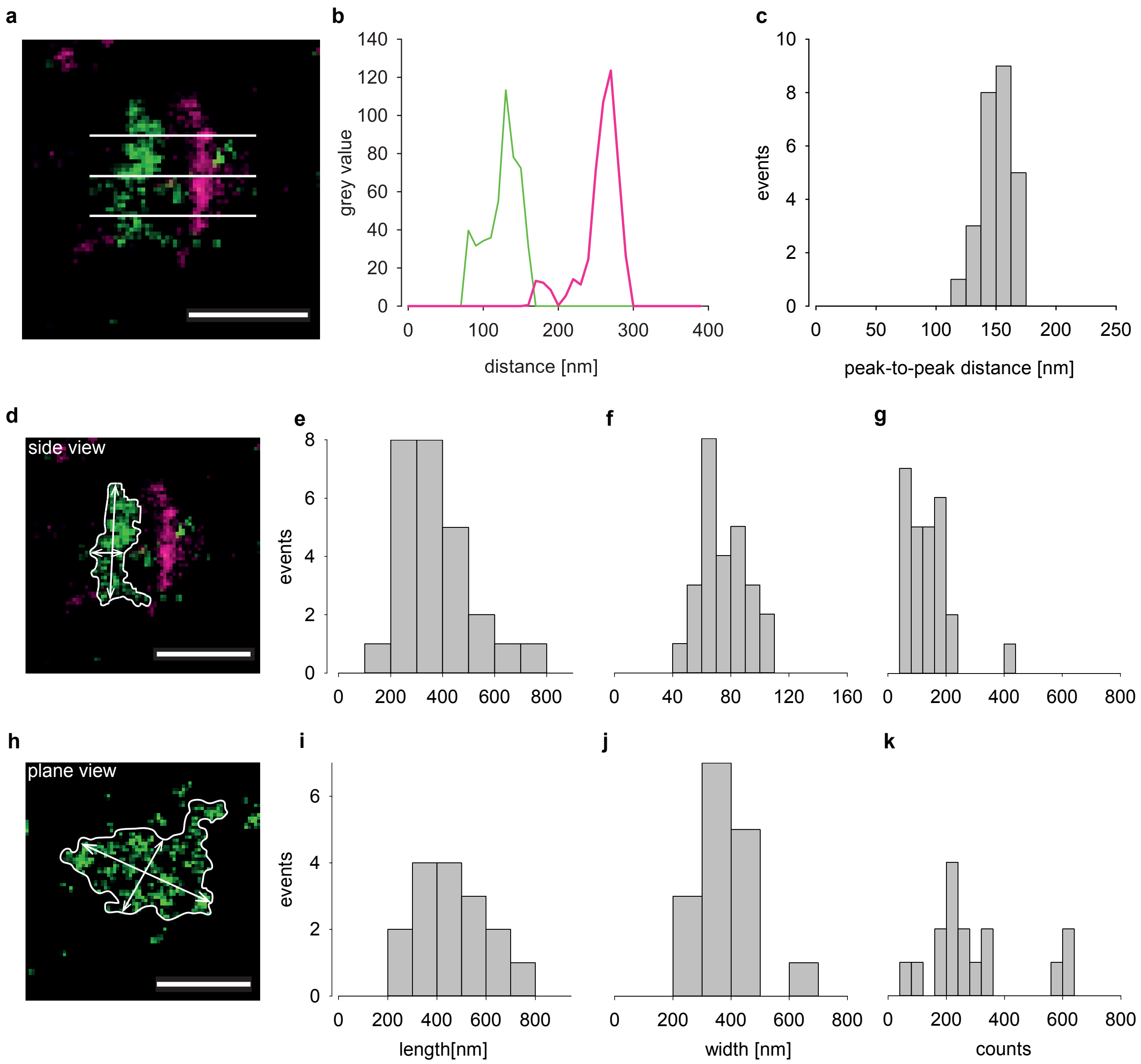


**Supplemental Fig. S1. Peak-to-peak-distance, Bassoon cluster length, width and localization counts in synaptic contacts in side view and in plane view.** (**a**) A representative image of an individual synaptic contact in side view with presynaptic Bassoon in green and Homer1 in magenta. White lines depict positions used for peak-to-peak measurements between pre-and postsynaptic protein clusters. (**b**) Grey values plotted against distance for peak-to-peak measurements between Bassoon (green) and Homer1 (magenta, measurement corresponds to central line in **a**). (**c**) Histogram of peak-to-peak distances for 26 individual synaptic contacts in one slice. (**d**) Representative *d*STORM image of an individual MF synapse in side view, Bassoon (green), Homer 1 (magenta). White line outlines Bassoon area. Arrows illustrate measurements of length and width. Histograms of length (**e**), width (**f**) and localization counts (**g**) of 26 individual Bassoon clusters in side view from one mouse. (**h**) Representative *d*STORM image of an individual MF synapse in plane view, Bassoon (green; Homer 1 not shown for display purposes) and illustration of analysis as in (**d**). Histograms of length (**i**), width (**j**) and localization counts (**k**) of 16 individual Bassoon clusters in plane view from the same mouse as in (**e**)-(**g**). Scale bars in (**a**), (**d**) and (**h**) 300 nm.

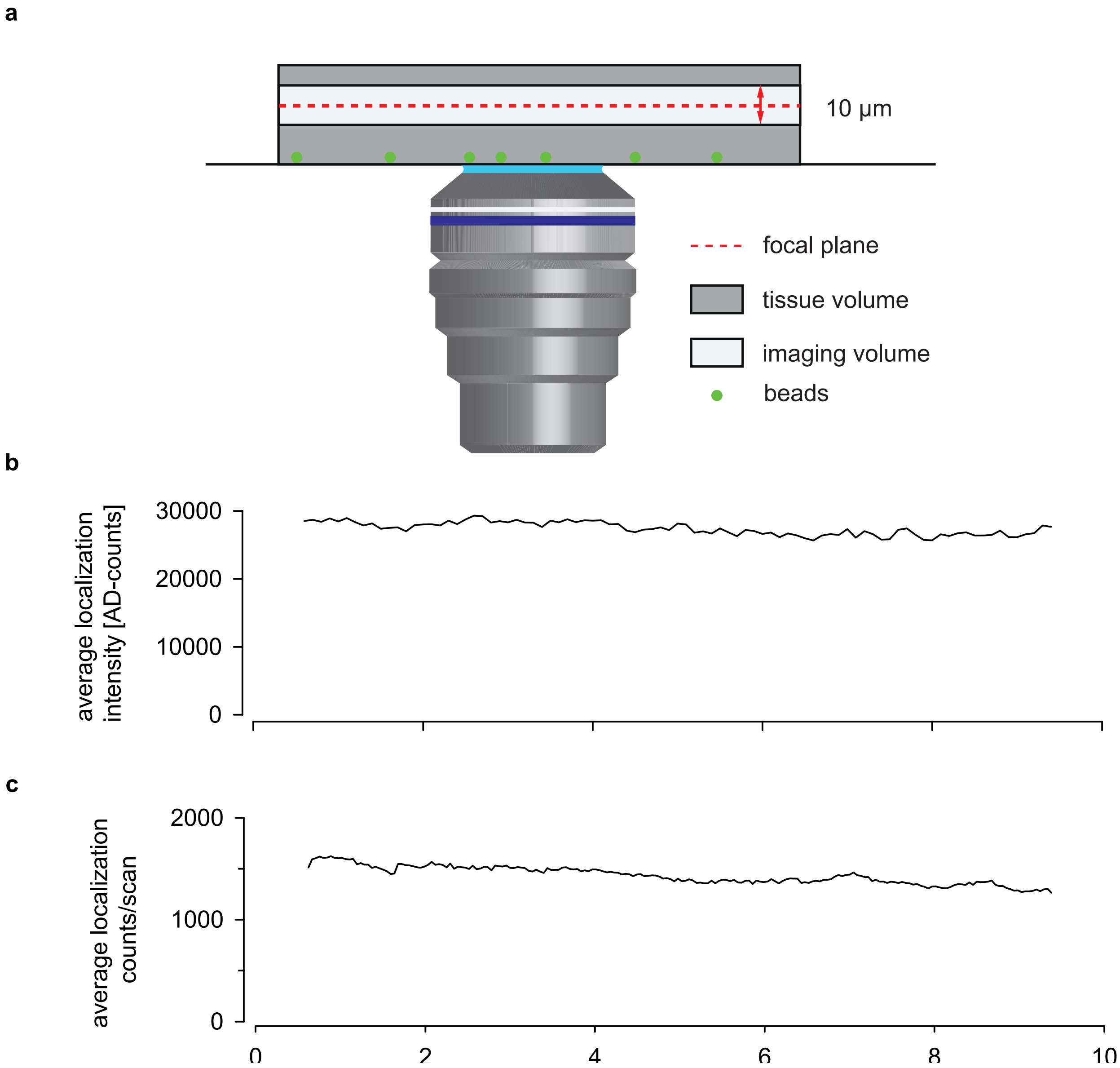


**Supplemental Fig. S2. Setup for continuous axial scanning and *en bloc* 3D imaging in thick tissue slices.** (**a**) Schematic illustration of the setup with a 25 µm-thick brain section on a fluorescent beads (green) -coated coverslip (not to scale). The relative position of the focal plane (red dotted line) is continuously stepped up and down (red arrow) through the region of interest (white). Average localization intensity in one scan (**b**) and average localization counts per scan (**c**) of all axial scans in one image are shown relative to focal position for the region of interest (see methods). Continuous axial scanning results in homogeneous distributions of AD-counts and density of localizations.

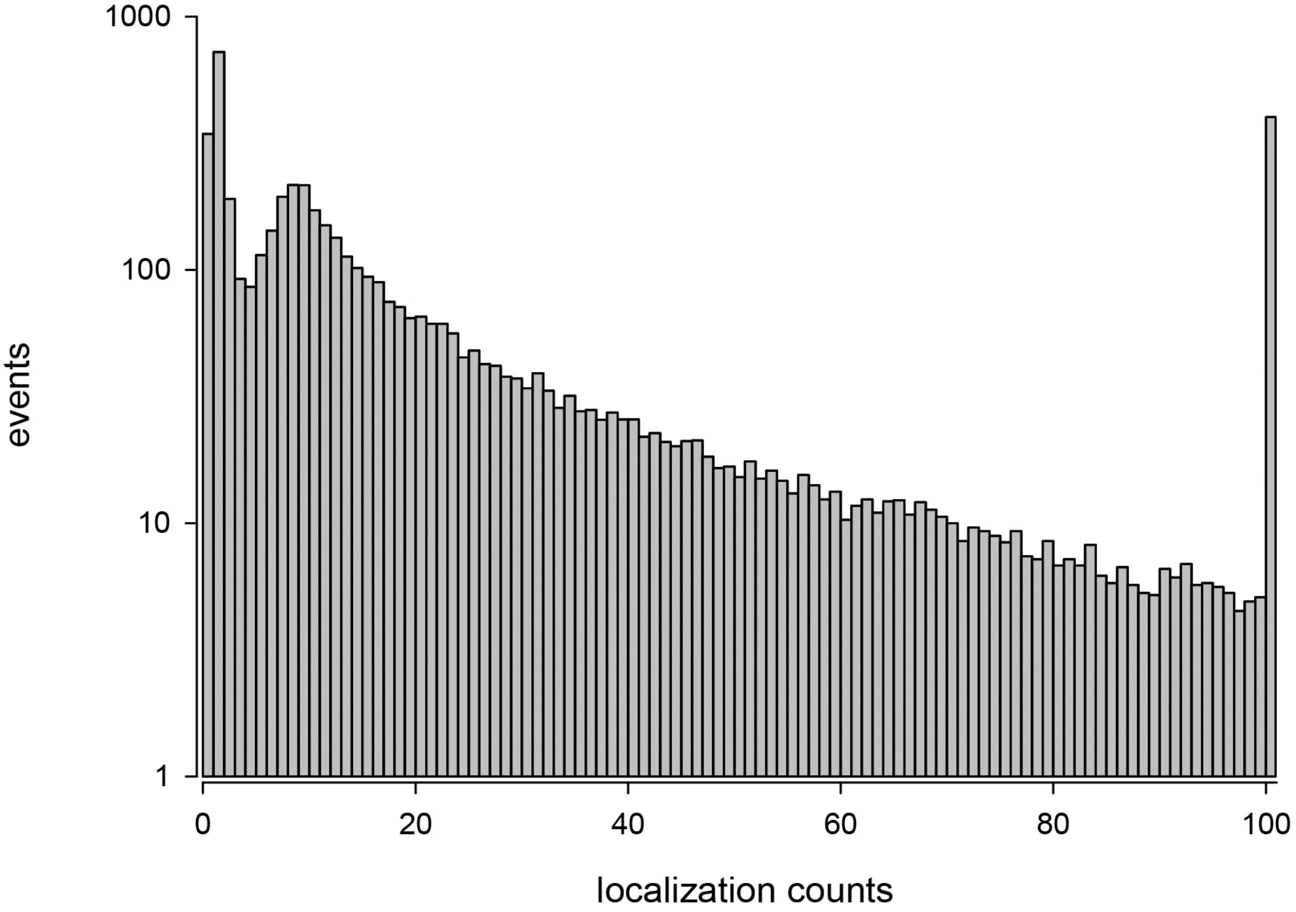


**Supplemental Fig. S3. Histogramm of localization counts per cluster in non-filtered data.** A cut-off of 8 localizations was used for filtered data.

| bouton- number | bouton-volume [µm$^3$] | number of clusters | cluster volume median [µm$^3$] | cluster volume range [µm$^3$] |
|---|---|---|---|---|
| bouton 1 | 20.81 | 15 | 0.0067 | 0.0026-0.0128 |
| bouton 2 | 13.84 | 17 | 0.00877 | 0.0021-0.134 |
| bouton 3 | 13.56 | 9 | 0.0152 | 0.0023-0.0568 |
| bouton 4 | 7.94 | 11 | 0.0086 | 0.0026-0.0481 |
| bouton 5 | 8.67 | 4 | 0.0250 | 0.0020-0.0478 |
| bouton 6 | 3.28 | 1 | 0.0219 | 0 |
| bouton 7 | 2.88 | 1 | 0.0283 | 0 |
| bouton8 | 2.65 | 1 | 0.0041 | 0 |
| bouton 9 | 2.33 | 5 | 0.0204 | 0.0033-0.108 |
| bouton 10 | 24.34 | 33 | 0.0211 | 0.0019-0.177 |
| bouton 11 | 2.59 | 7 | 0.0177 | 0.0045-0.264 |
| bouton 12 | 2.69 | 2 | 0.0433 | 0.0354-0.0512 |
| bouton 13 | 4.30 | 8 | 0.110 | 0.0022-0.279 |
| bouton 14 | 3.18 | 7 | 0.0052 | 0.0004-0.0149 |
| bouton 15 | 1.33 | 1 | 0.0054 | 0 |
| bouton 16 | 3.85 | 4 | 0.0173 | 0.0069-0.112 |
| bouton 17 | 4.36 | 7 | 0.0406 | 0.0181-0.139 |
| bouton 18 | 2.91 | 1 | 0.0620 | 0 |
| bouton 19 | 3.79 | 1 | 0.105 | 0 |
| bouton 20 | 32.18 | 45 | 0.0134 | 0.0018-0.167 |
| bouton 21 | 3.08 | 1 | 0.0100 | 0 |

**Supplemental Table S1. Characteristics of Bassoon clusters in individual mossy fiber boutons.**

**Supplemental Video 1. 2-color 3D *d*STORM volumetric presentation of a single synaptic contact with Bassoon (green) and Homer 1 (magenta).**

**Supplemental Video 2. Sequential 3D *d*STORM volumetric presentation of Bassoon clusters and mEGFP in mossy fiber tract of a Thy1-mEGFP (Lsi1) mouse.** Bassoon clusters are color-coded according to cluster size from blue to red, mEGFP in axons and boutons in cyan.
